# TopOmics: Topic Modelling for All Omics

**DOI:** 10.64898/2026.05.26.727810

**Authors:** Federico Caretti, Nour El Kazwini, Guido Sanguinetti

## Abstract

Topic models have emerged as a popular paradigm to analyse and interpret complex single-cell and spatial data. Yet, current implementations are usually data-type specific and rely on different modelling and estimation approaches, hindering usability and interoperability. In this work we introduce TopOmics, a library to perform efficient and flexible topic modeling with any combination of -omics data at scale. The framework leverages standard libraries of the Python ecosystem, guaranteeing seamless integration with existing pipelines, and shows competitive performance against state-of-the-art methods while preserving interpretability. We provide several examples of TopOmics on diverse data sets, including a novel topic model for spatial multi-omic data, and an analysis of a very large VisiumHD data set.

## 1 Introduction

Biological complexity arises from interactions of many molecular factors at varying spatial and temporal scales. Major advances in sequencing technologies have enabled the direct measurement of many such molecular players [1, 2, 3], as well as the quantification of their spatial variability at the single cell or near-single-cell resolution [4, 5]. Coupled with the diminishing costs of sequencing itself, these advances have led to a proliferation of highly complex, high-dimensional data sets profiling millions of cells across different tissues and biological processes.

Such growing complexity and scale of data poses a direct challenge to the modelling community: summarising such wealth of data efficiently and informatively calls for scalable, accurate and interpretable unsupervised learning techniques. Proposed solutions range from deep learning-based methods [6, 7, 8] to multi-tasking factor analysis [9], striking different balances along the expressivity-interpretability trade-off.

In this context, topic models have emerged as a powerful framework that strikes an attractive balance between flexibility and interpretability: they can give rise to complex multimodal data distributions, they can easily be extended to encompass nonlinear encoding mechanisms, yet they maintain an interpretable-by-design nature by associating each latent direction with a “topic” which recapitulates essential directions of variation within the data. Initially developed in the context of natural language processing and automated annotation [10, 11], they have been successfully deployed to model single-cell transcriptomic [12, 13], epigenomic [14], multi-omic [15, 16] and spatial transcriptomic data [17, 18]. While the results presented in these papers support a prominent role for topic models in the single-cell analysis toolbox, their implementation is usually data-type specific, with different noise models, different priors and different estimation algorithms used in each separate package.

In this paper, we present TopOmics, a modular framework for topic modelling in single-cell and spatial ‘omics. Built around an amortised variational inference engine which can easily scale to large data sets, TopOmics provides the user with a number of likelihood options which enable flexible modelling of different types of data. As a demonstration of the ease of application of TopOmics, we present results on spatial multi-omics and single-cell triple ‘omics data; in both cases, TopOmics is the first topic modelling framework that can handle this data type.

## 2 Results

### 2.1 Model description and computational efficiency

TopOmics is a flexible framework for topic modelling of any combination of omics, independently of the sequencing technology used. Topic models were originally developed in the context of automatic document annotation in natural language processing: in the latent Dirichlet allocation (LDA) model [10], each word in a document is assigned to a specific topic, while the document itself is characterised by a distribution of topics which categorise it (see Figure 1 a). In a biological context, topics are identified with biological processes which are happening in individual cells (which correspond to documents in the original NLP application), while genes and other molecular entities such as chromatin regions play the role of words (see Figure 1 b). In multi-omics, we have multiple observation modalities for each cell (which are treated as multiple versions of the same document), and we enable spatial correlations between cells (which are dealt with as correlated documents, where a correlation structure is provided).

**Figure 1:**
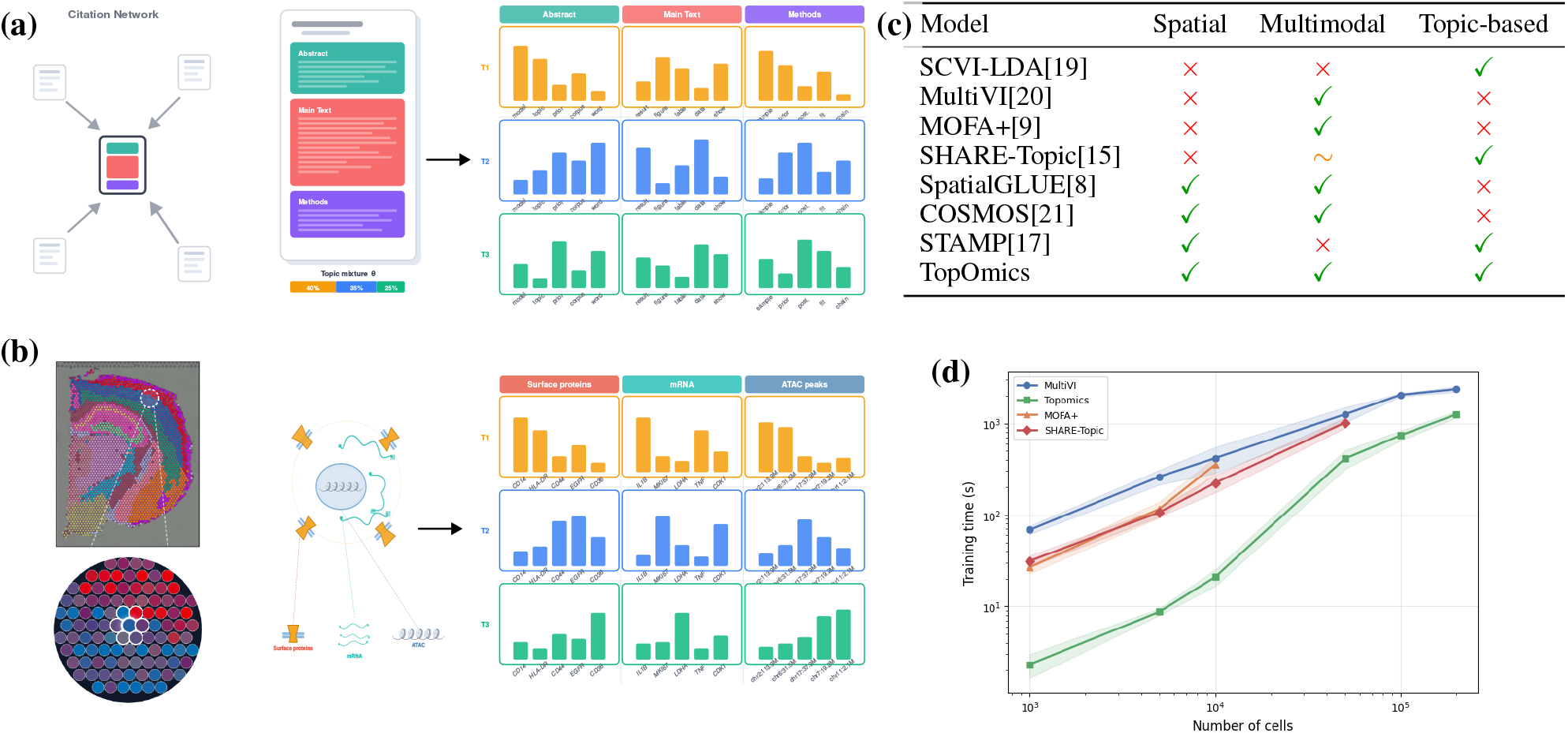
(a) In traditional topic modeling, word counts in a document are modeled through the contribution of different topics. Extensions of topic models can also be applied to networks of documents, such as citation networks[22], and can assume different words distributions for the documents’ sections. (b) In the application of topic models to -omics data, cells or spots are the “documents” and the observed modalities take the role of the different sections of the documents. Instead of citation networks, spatial information can be incorporated in the form of a spatial neighborhood graph (c) Comparison with competitive methods used for comparison and their ranges of applicability. (d) Scaling times of TopOmics, SHARE-Topic, MultiVI and MOFA+ on synthetic RNA+ATAC datasets with variable numbers of cells. We were unable to run MOFA+ and SHARE-Topic on 10^5^ or more cells; more details on the experimental setup in Suppl. Sect. D and on our hardware in Sec.6.

The implementation of TopOmics is based on a core inference engine based on an amortized[11] version of LDA. As a prior distribution over topics, we implemented both the standard Dirichlet prior, and the Horseshoe prior [23], which encourages sparsity. The different biological modalities correspond to different observation models, with pre-implemented models for RNA, proteins and chromatin accessibility. This gives TopOmics considerable flexibility making it adaptable to many different types of modern ‘omics experiments (see Figure 1 c).

The amortized implementation requires a GPU for fast inference, but provides great scalability both in the number of features and in the cells. Figure 1d shows the runtime of TopOmics, SHARE-Topic, MultiVI and MOFA+ on synthetic datasets of varying size (see Methods for details of how these data sets were generated). We apply early stopping to MultiVI, TopOmics and SHARE-Topic if the validation loss (ELBO for the first two, log-likelihood for SHARE-Topic) does not decrease for 50 epochs, while using the default greedy strategy for MOFA+. Not only TopOmics shows the fastest runtime, but the minibatching scheme avoids loading all the samples into the memory at once, allowing it to scale to numbers of cells that are prohibitive for MOFA+ and SHARE-Topic.

### 2.2 TopOmics recovers tissue architecture from spatial multi-omic data

We test TopOmics on a dataset of spatial epigenome-transcriptome co-profiling of the coronal section of the mouse brain from [24]. On this dataset, we benchmark TopOmics against four competing methods: SpatialGlue[8] and COSMOS[21], which are designed for spatial multi-omic data, MultiVI[20] and MOFA+[9] that neglect the spatial information. The ground truth annotation was performed in [21] by using the Allen Brain Atlas (https://mouse.brain-map.org/static/atlas) as a reference for the annotation of the anatomical regions.

We apply two variants of TopOmics, testing both the Dirichlet and Horseshoe priors over the topics. For models that support spatial information, the neighborhood graph was built considering only the first neighbors on a square grid. We ran all models with 10 and 20 latent dimensions, and the latent space classification was performed using the KMeans clustering algorithm with *k* = 8 (the number of clusters in the annotation) and *k* = 15. We evaluate quantitatively the quality of the results of the various methods by computing the accuracy of spot classification and the normalised mutual information (NMI) against the ground truth annotation, as well as the Moran I score which measures spatial consistency of an annotation. Only annotated spots were considered during the evaluation of the computed metrics.

First, we observe that the TopOmics latent representations qualitatively strike an excellent balance between capturing the anatomy of the mouse brain while also aggregating information from the neighbors and denoising the latent representations. MOFA+ can distinguish some anatomical features, such as the lateral ventricle (VL), but struggles to separate the other anatomical regions (Fig.2a) and has the lowest performance in both settings. COSMOS (Fig.2d) performs well in general and is particularly strong on the Moran I metric, indicating high spatial consistency, but at the cost of introducing some spatial artifacts from the local neighborhood aggregation. MultiVI and SpatialGLUE (Figs.2b2c) perform generally well but struggle to separate the different cortical layers; the latter boosts a higher Moran’s I score thanks to the spatial aggregation strategy. TopOmics is capable of correctly distinguishing the different cortical layers without introducing noticeable artifacts, and performs best across almost all metrics.

**Figure 2:**
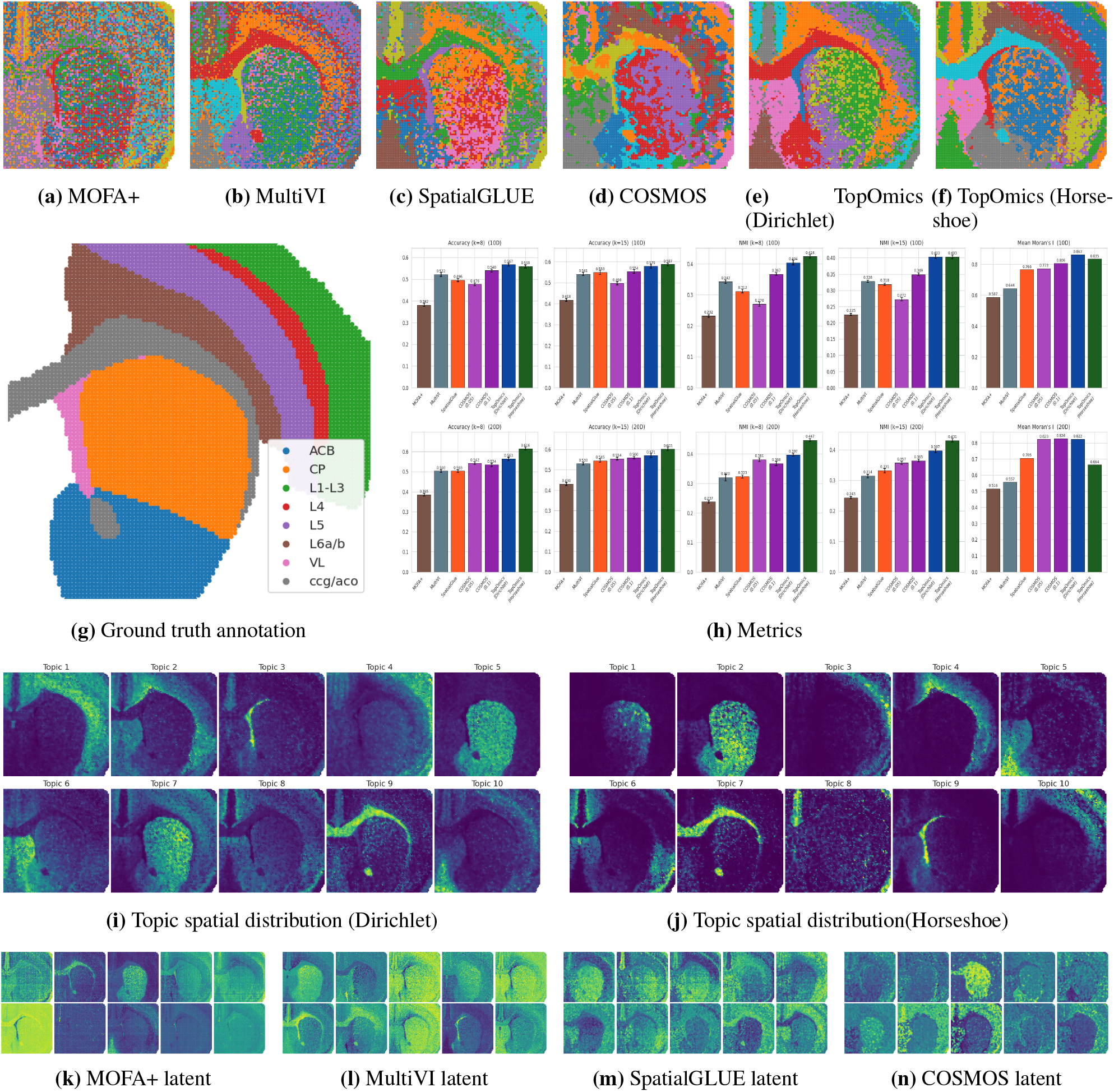
2(a-f) K-means clustering results (k = 15) for all compared models using a latent dimension (number of topics for TopOmics) of D = 20. (g) Ground-truth annotations from [21]. (h) Quantitative evaluation using a kMeans classifier, NMI, and Moran’s I across varying numbers of clusters (k = 8 and k = 15) and latent dimensions (D = 10 and D = 20). (i–n) Spatial distributions of topics/latent factors learned with D = 10.

TopOmics also provides more interpretable latent factors than the competing methods, as shown in Figs.2i2j. In many cases, a single topic is strongly associated with a specific anatomical annotation, both with the Dirichlet and Horseshoe priors. By contrast, Figs. 2k2l2m2n show that all the other methods produce significantly less diverse latent representations which are not directly interpretable.

In summary, this analysis demonstrates that TopOmics provides a powerful platform for modelling complex spatial multi-omic data, achieving state-of-the-art anatomical reconstruction and providing full interpretability.

### 2.3 TopOmics scales efficiently to spatial cell-level resolution datasets

We demonstrate the flexibility and scalability of the TopOmics framework by applying it on two VisiumHD samples of colorectal cancer[5] from two different patients, here reported as P1 and P2. Cells are segmented using the QuPath[25] software and default parameters; nucleus detection is aided by the Hematoxylin channel, obtained via color deconvolution of the H&E image, and cytoplasm detection is performed by expanding the cells around the nucleus. Two different ground-truth annotations are provided for each sample [5], one coarse-grained (10 labels, as reported in Figs. 3a 3b) and one fine-grained (39 and 41 labels respectively). The cells labeled as Unknown were included in the training set but not considered during the evaluation of the metrics in Figs.3g3h.

**Figure 3:**
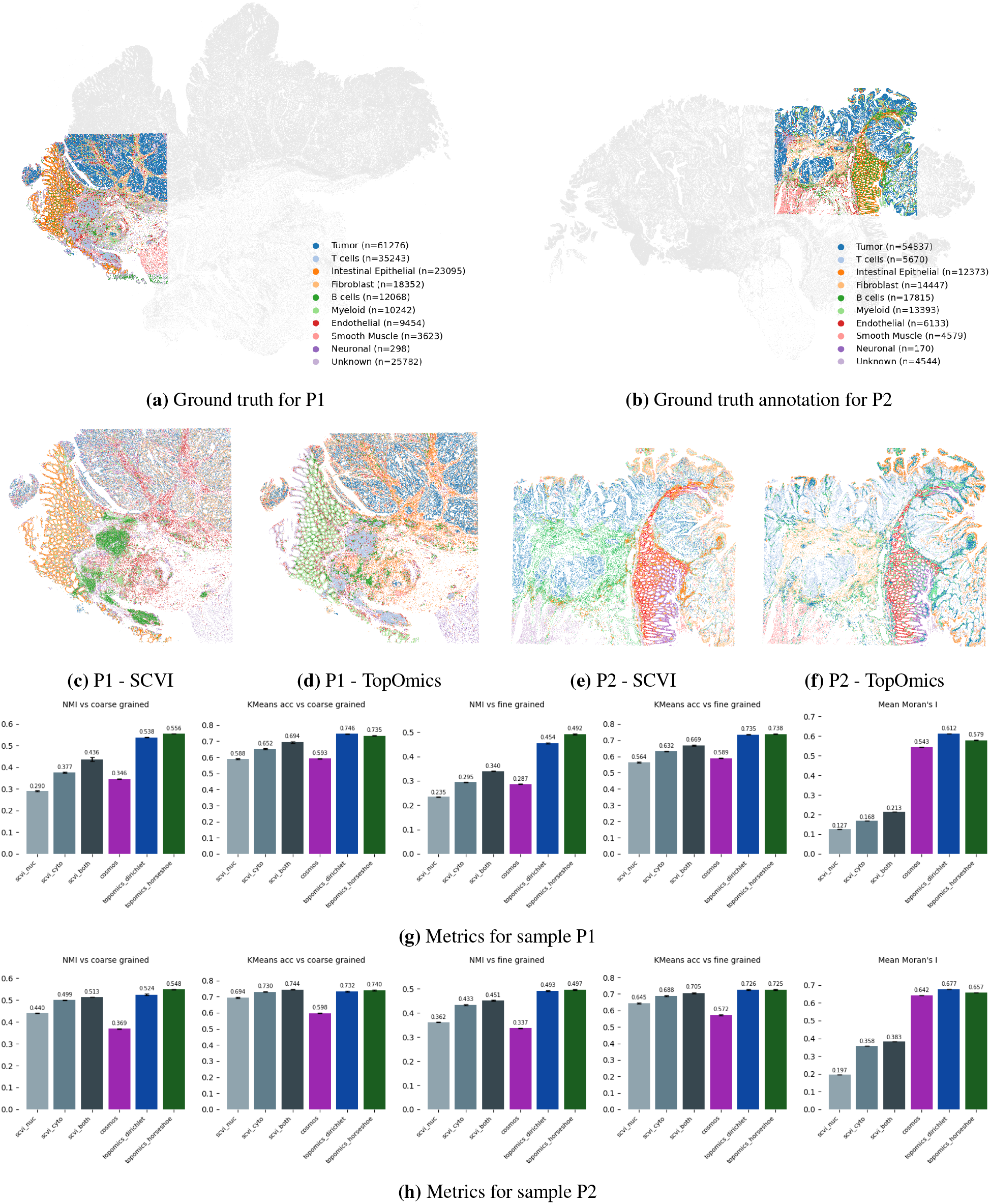
3a3b ground truth annotations for the two samples from Patient 1 (P1) and Patient 2 (P2) respectively. 3c3d(3e3f): latent representations generated by scVI (nucleus and cytoplasm combined) and TopOmics (Horseshoe prior) on samples P1 (P2 respectively). 3g(3h): comparison metrics for scVI, COSMOS and TopOmics on sample P1 (P2)

The main challenge in this dataset is represented by the high number of cells in the sample, which leads to severe compute and memory problems for methods that do not rely on mini-batching. We run TopOmics, both with Dirichlet and Horseshoe prior, treating cytoplasmic and nuclear RNA as two separate modalities. We compare the performance of TopOmics against three different variants of SCVI[6] (considering nucleus RNA or cytoplasmic RNA only, or both) and against COSMOS[21] trained similarly to TopOmics. We were unable to run either STAMP[17] or SpatialGLUE[8] as the very large number of cells led to memory issues.

Figs.3g3h report the Normalized Mutual Information and performance of a KMeans classifier for all models, on both datasets and against both the coarse-grained and the more detailed labels, and Moran’s I score on both datasets. Particularly on the P1 data, both variants of TopOmics outperform the competitors by a notable margin. The difference between the scVI variants shows that cytoplasmic RNA is significantly more reliable for cell type classification, but there is an improvement in using both. We again notice that COSMOS achieves a very high spatial correlation at the cost of accuracy on the other metrics, while both TopOmics variants perform similarly and are the only models able to balance between accuracy and spatial smoothness in the latent space projections.

### 2.4 TopOmics scales to any number of modalities

While most multimodal single-cell ‘omic data sets are limited to two modalities (generally transcriptomic and chromatin accessibility), new technologies are emerging that can measure at scale more than two modalities in individual cells. To showcase the flexibility of TopOmics, we examined a TEA-seq dataset of neutrophil-depleted PBMCs[2]. TEA-seq simultaneously measures transcriptomes, chromatin accessibility and a panel of surface proteins in every cell. A major challenge in this type of data is the huge heterogeneity in the number of features and informativeness among the modalities: although surface proteins are the best modality for cell type classification, they are by far the modality with the lowest number of features, while the opposite holds true for chromatin, as shown in the metrics in Fig.4g.

**Figure 4:**
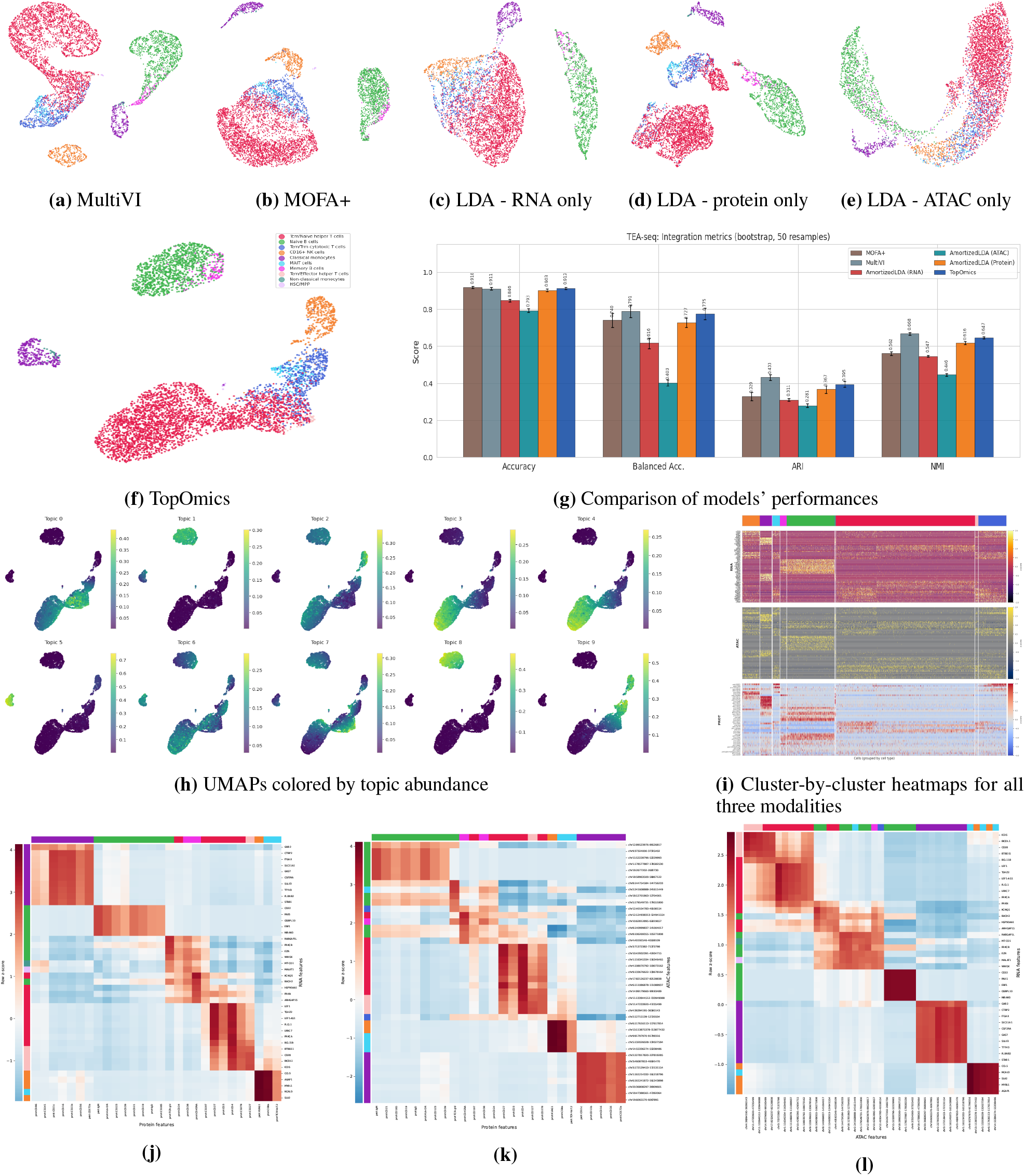
UMAPs of the latent representations generated by different methods: MultiVI (4a), MOFA+ (4b), TopOmics (4f) and unimodal topic models of the class AmortizedLDA from SCVI-tools[19] (RNA (4c), proteins (4d) and chromatin accessibility peaks (4e) respectively). (4g): benchmark of the latent representation generated by TopOmics against those of the aforementioned models. (4h): activation of each topic in the latent space. (4i): For every cluster, some of the most relevant differentially expressed features are highlighted, according to their inferred distribution means. (4j), (4k), (4l): heatmaps of feature-feature interactions.

We first annotated the cell types using CellTypist[26] and then compared the performance of TopOmics to MOFA+[9], MultiVI[20] and single-modality versions of the AmortizedLDA class from SCVI-tools[19].

All methods yielded qualitatively similar visualisations which highlight the presence of numerous cell clusters(Figure 4 a-f). Quantitatively, multimodal methods perform better than single-modality methods, indicating that each modality contributes some independent information towards cell type detection (Figure 4g). TopOmics achieves this while retaining interpretable topic-level representations (Fig.4h).

TopOmics allows a range of possible downstream analyses, which can inform about the relevance of individual features towards the cell classification, and about the interactions between different modalities. Figure 4i shows an example heatmap highlighting the features which are inferred to have the highest relevance towards cell type classification. These recapitulate known biological markers such as the CD16 protein for CD16+ NK cells and the CD14 protein for classical and non-classical Monocytes[27].

Fig.4 (j-l) show instead interactions among features, which can be used to help formulate mechanistic hypotheses. For example, Figure 4l details pairs of strongly interacting chromatin and RNA features, detailing candidate instances of epigenetic regulation of the relative target genes. As the heatmaps in Fig.4l were generated using the whole dataset, the clusters significantly correlate with the cell types, whose colors are indicated at the borders. See AppendixD for details of how these figures were generated.

## 3 Discussion

The increasing scale and complexity of biomolecular data sets calls for effective and efficient methodologies to analyse them. TopOmics is a comprehensive framework for interpretable modelling of data from many modern single-cell or spatial ‘omics technologies. Using an efficient amortized implementation of topic modeling, it can scale to very large multi-omic and spatial multi-omic data sets. In this work, we showcased state-of-the-art performance on datasets at different resolutions, with or without spatial information, and with different amount of modalities. This includes novel implementations of topic modelling for spatial multi-omics and Visium HD data sets.

TopOmics is interpretable-by-design, as it explains the data distribution as a generalised linear combination of topics, which themselves depend (non-linearly) on the data in an auto-encoding fashion. Compared to fully black-box architectures such as SCVI[6] and MultiVI[20], the generalised linear structure of the decoding map limits the theoretical expressivity of TopOmics. In practice, we show that TopOmics achieves state-of-the-art performance on multiple benchmarks and across different metrics; its lower flexibility seems therefore to be a limited price to pay for the interpretability afforded by the topic modelling formulation.

TopOmics does carry some notable limitations. The amortized inference engine, while enabling scalability to very large datasets, in practice requires a GPU. Moreover, replacing the black-box decoder with an interpretable generative model introduces the need to specify additional hyperparameters beyond the number of topics (latent variables) K, such as the concentration parameters of the Dirichlet distribution. While we provide reasonable defaults, spatial applications in particular require the user to tune the spatial aggregation parameters depending on the downstream task, which can range from tissue segmentation to cell type classification.

With these tradeoffs in mind, TopOmics is designed to be accessible to all users, with automatic hyperparameter selection based on the input data, and full compatibility with popular single-cell analysis frameworks such as scverse. The implementation remains flexible to support use cases beyond those discussed in this paper, such as incorporating prior knowledge to construct cell graphs or applying the model on top of external batch correction methods. This flexibility will also in principle enable easy adaptation to yet-to-be-developed sequencing-based single cell technologies, such as high resolution spatial multiomics.

## 4 Methods

TopOmics is an extension of topic models to single-cell multiomics data, optionally incorporating spatial information. A conceptual illustration of the TopOmics model is shown in Figure 6.

**Figure 5:**
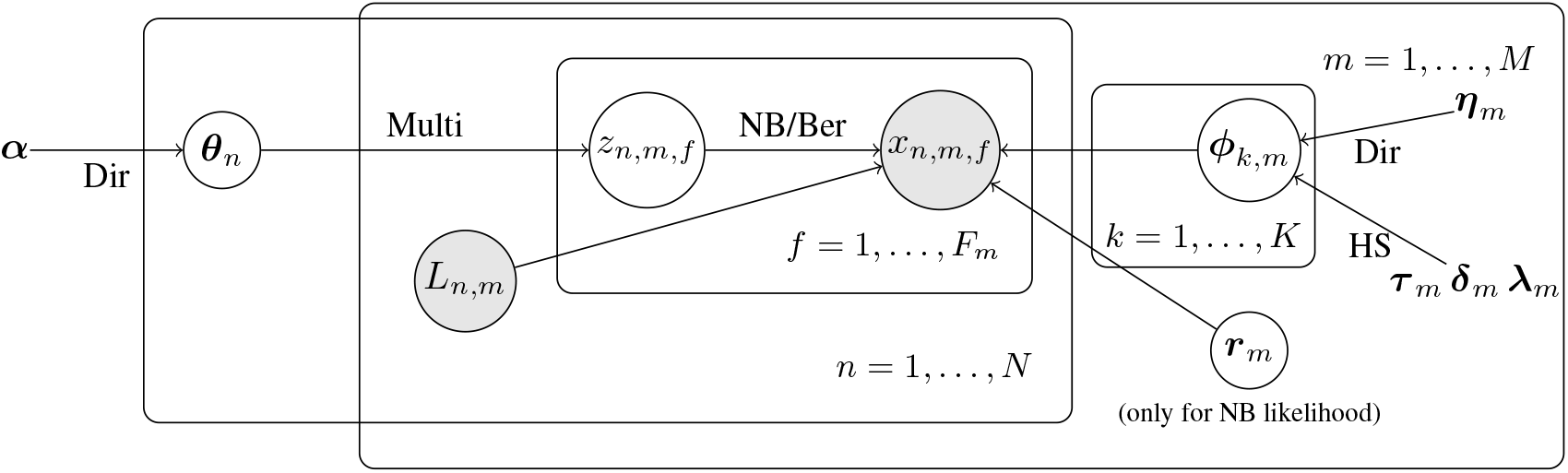
The generative model of TopOmics. Each cell is assigned a distribution over topics ***θ***_*n*_, drawn from a Dirichlet prior with concentration ***α***. For each modality *m*, a topic-feature distribution *ϕ*_*k,m*_ is learned under either a Dirichlet (Dir) or Horseshoe (HS) prior. Topic assignments *z*_*n,m,f*_ are drawn from a Multinomial distribution and observations *x*_*n,m,f*_ are generated via a Negative Binomial, which also needs a dispersion parameter ***r*** or Bernoulli likelihood, depending on the data type (or other options mentioned in Suppl.A). The library size *L*_*mn*_ is removed from the inputs by normalizing the total sum of reads per data point, and injected in the model as a parameter of the likelihood function. Shaded nodes denote observed variables.

**Figure 6:**
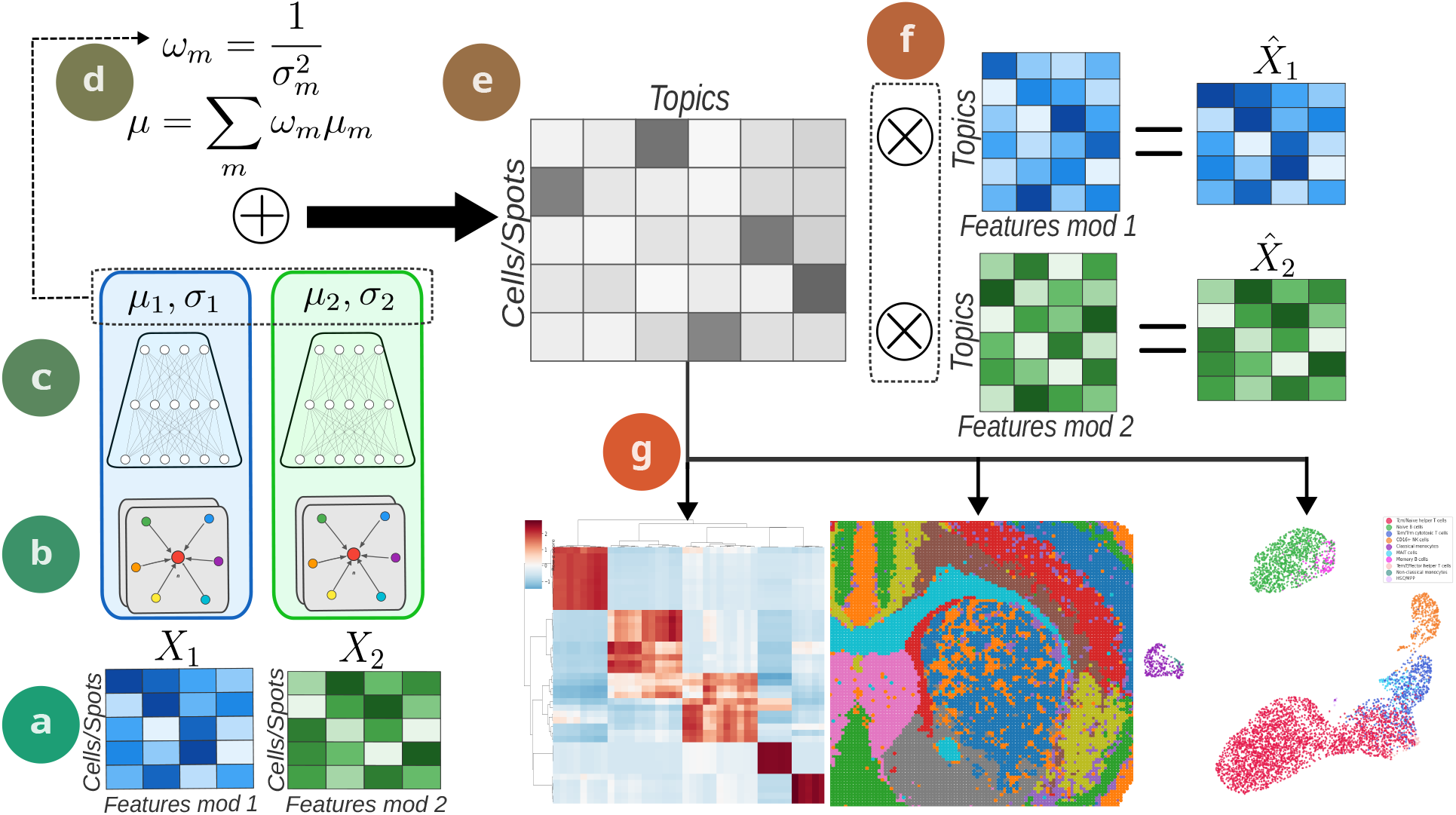
Algorithm overview. **(a)**: data matrices representing the input data corresponding to different modalities. Inputs are normalized per-cell (or spot) so that the encodings are independent of the library size of each point. **(b)** In the presence of spatial information, TopOmics builds neighborhood graphs and first applies graph attention layers. **(c)** The standard encoder used by TopOmics for every modality is a fully connected 2-layers neural network that outputs the parameters *µ* and *σ* for every modality. **(d)** TopOmics then combines the information coming from the modalities through a weighted Mixture-of-Experts; by default, the weight assigned to each modality is cell-dependent and corresponding to the inverse of the variance provided by the encoder. **(e)** By applying a SoftMax function to the mixture, the latent representation of each cell or spot sums to one. This is the standard implementation of the amortized approximation of the Dirichlet distribution, as in [11]. **(f)** The matrix product of the latent representations and the topics-features parameters returns the average values of the features, under the possible likelihood functions. The model is trained to maximize the likelihood that the data come from the decoded distribution. **(g)** Some of the downstream tasks that TopOmics can perform after training include detection of normalized feature-feature interactions, segmentation of spatial domains and clustering.

A more precise description of the generative process of TopOmics is illustrated using the language of probabilistic graphical models in Figure 5. The observables *x*_*mn*_ consist of *M* vectors (of different lengths *F*_*m*_ *m* = 1, … , *M*) per each of *N* cells (or spots). These represent the observed modalities (e.g., transcriptomes, chromatin accessibility, nuclear RNA, etc); each cell has an additional observed variable *L*_*m,n*_ which is generally derived from the *x*_*mn*_ observable and used for normalisation purposes (e.g., for transcriptomic observations, *L*_*m,n*_ could be the total reads mapped in the cell). Each feature in each modality in each cell/spot is associated to a discrete latent variable *z*_*nmf*_ taking one of *K* topic values; the probability of taking each of the values is specified by a cell/ spot level distribution ***θ***_*n*_ which is given a Dirichlet prior distribution with concentration parameter ***α*** (a hyperparameter set by default to a constant vector ***α*** = *α*1). Lower values of ***α*** push the model towards sparse and peaked topic distributions, while higher values nudge the model to use more, less distinct topics to explain the observations. The topic-features distribution ***ϕ***_*m*_ indicates the distribution of the features of a given modality given the topics: *x*_*n,m,f*_ is sampled from the distribution corresponding to the variable *z*_*n,m,f*_ . See e.g. [10] for a thorough explanation of topic models.

In formulas, the generative process is therefore the following:

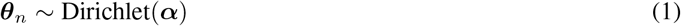

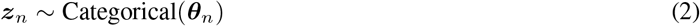

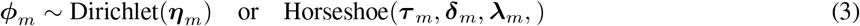

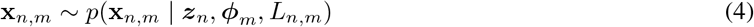

### 4.1 Posterior approximation

TopOmics is an *amortized* model, meaning that the posterior distribution has a specified parametric form with parameters depending on the input data through a neural network. We explain the procedure for a single modality *m*, following [11]; multiple modalities are then combined according to the strategy described in subsection 4.4. For each cell/ spot *n*, the approximate posterior over the topic distribution ***θ***_***n***_ is obtained as the softmax transformation of a spherical Gaussian with parameters ***µ***_*n*_(**x**_*n*_), ***σ***_*n*_(**x**_*n*_) ∈ ℝ^*K*^ , with *K* being the number of topics.

The topic-features parameters ***ϕ***_*m*_ undergo a similar procedure in the case of the Dirichlet prior, for which the same analytic approximation of the KL divergence is available. In the case of the Horseshoe prior, since no formula is available, the KL divergence is estimated via Monte Carlo sampling.

We perform amortized variational inference, optimizing the evidence lower bound (ELBO):

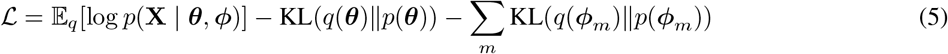

in which the likelihood term (first) pushes the model to fit the data, while the KL regularization terms penalize deviations of the approximate posteriors from their respective priors, preventing overfitting. See [11] and the implementation in [19] for details of the optimisation procedure.

The encoding network is, by default, a standard 2 layers fully connected neural network, preceded by some Graph Convolution layers in the case of spatial datasets. The input is the log of the data, normalized per-cell by library size. Input data are split into training and test set, and overfitting is prevented by early stopping depending on the test loss.

### 4.2 Likelihood Functions

TopOmics supports four likelihood functions to accommodate different data modalities.

For overdispersed count data (RNA-seq and proteins), we use a Negative Binomial likelihood:

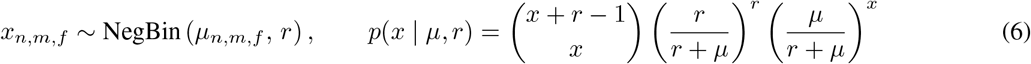

where *µ*_*n,m,f*_ = *L*_*n,m*_ ∑ _*k*_ *θ*_*n,k*_ *ϕ*_*k,m,f*_ scales expected expression by the library size *L*_*n,m*_, and *r >* 0 is a learnable dispersion parameter.

For binary presence/absence data such as ATAC-seq peaks or methylation sites, the Bernoulli likelihood is the natural choice:

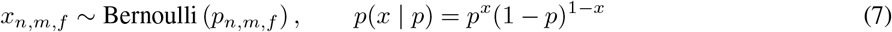

where 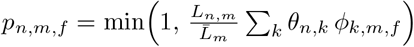 and 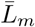 is the median library size for modality *m*, ensuring *p*_*n,m,f*_ ∈ [0, 1].

Furthermore, for modalities with continuous or non-integer values, for instance after external batch correction with Harmony[28], we support a Gaussian likelihood, corresponding to Gaussian LDA[29]. TopOmics also supports the Multinomial likelihood that is the standard in topic models. More details in Suppl.A.

### 4.3 Topic-Feature Priors

We implement two prior choices for the topic-feature distributions 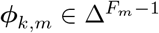.

The default is a symmetric Dirichlet prior, shared across all topics *k* of a given modality *m*:

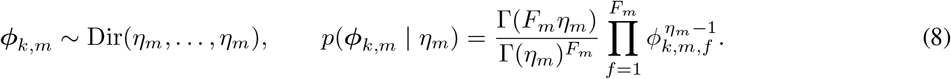

The concentration *η*_*m*_ *>* 0 controls sparsity: *η*_*m*_ *<* 1 pushes mass toward the corners of the simplex (sparse topics, few features per topic), *η*_*m*_ = 1 gives a uniform prior on the simplex, and *η*_*m*_ *>* 1 concentrates mass near the centroid (dense topics). By default we set *η*_*m*_ = 1*/K*. In practice, because the Dirichlet distribution does not support any exact reparameterization trick, we apply the same principle followed for the cell-topic distribution ***θ*** and approximate the Dirichlet distribution through the logistic-normal approximation.

The second option is the Horseshoe prior[23], which has been introduced in the context of topic models for spatial omics in STAMP[17] and is designed to force sparsity in the topic-feature parameters. The Horseshoe prior achieves this through a hierarchy of Half-Cauchy scales: a topic-level scale *τ*_*k,m*_ controls global shrinkage within each topic, a feature-level scale *δ*_*f,m*_ allows individual features to escape shrinkage across all topics, and a topic-feature scale *λ*_*k,f,m*_ permits specific associations to remain large even when the global scales are small.

Formally, for each modality *m*:

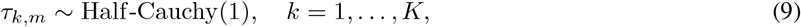

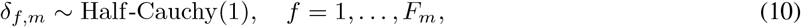

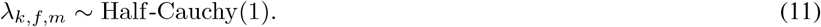

Then, a modality-specific parameter *c*_*m*_ is introduced to regularize the maximum value that the actual interaction parameter 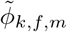 can assume, bounding it between 0 and 1:

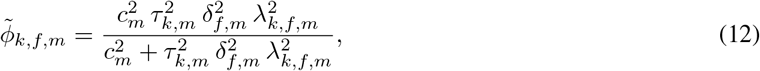

Where 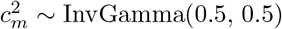.

Differently from the Dirichlet prior, for which an analytic formula for the KL divergence term is available, in the case of the Horseshoe prior the regularization term is estimated via Monte Carlo sampling.

Empirically, we observed that the increased topic sparsity helps balancing the smoothing introduced by the graph convolutions in the spatial applications, but the absence of an analytic formula for KL divergence hinders the stability of training if applied to non-spatial variants. For this reason, the Dirichlet prior remains the default choice in TopOmics for non-spatial applications.

### 4.4 Multimodal Integration

TopOmics employs a Mixture-of-Experts[30] architecture to infer a shared cell-topic distribution from multiple modalities. Each modality *m* has a dedicated encoder network *E*_*m*_ that maps the observed features of each cell or spot (indexed with *n*) to parameters of a Gaussian distribution in logit space:

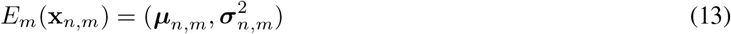

The modality-specific distributions are combined via weighted Gaussian mixture:

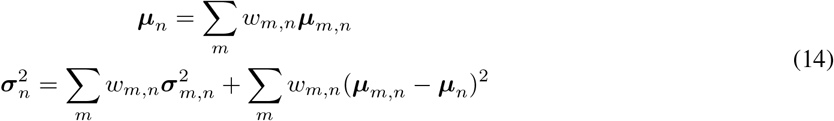

By default, the weights *w*_*m,n*_ are set to be inversely proportional to the variance of each modality, in order to downweight the modalities that are deemed unreliable by the encoders. Other possible choices for aggregation strategies are discussed in Suppl.A. The KL divergence term in Eq.5 with the prior coefficients *µ*_*k*_, *σ*_*k*_ is calculated w.r.t. the aggregated distribution; we show in Suppl.F that this choice implicitly nudges correlation across different modalities.

### 4.5 Spatial information aggregation

For spatially resolved datasets, we add to the inference network a graph neural encoder that propagates information between contiguous cells or spots. From the spatial coordinates, we construct a graph *G* = (*V, ℰ*) using the neighborhood information; in particular, for the VisiumHD dataset of Sec.2.3 we linked each cell with its 10 nearest neighbors, while for the lower resolution spatial dataset of Sec.2.2 in which the spots are distributed on a square grid we used the 4 nearest neighbors.

If such neighborhood graph is provided, TopOmics applies a few layers (2 by default) of Graph Attention layers[31]:

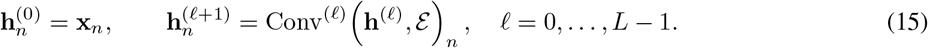

where Conv^(*ℓ*)^ denotes a GATv2 [32] convolution layer. For each edge (*i, j*) ∈ *ℰ*, the attention coefficient is computed as:

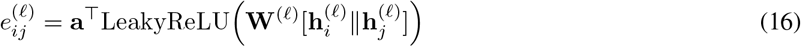

where **W**^(*ℓ*)^ is a learnable weight matrix, ∥ denotes concatenation, and **a** is a learnable attention vector. The coefficients are normalized across the neighborhood N (*i*) of node *i*:

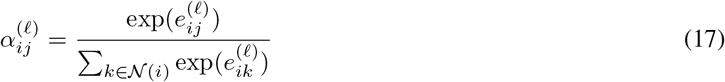

The output of the layer is then the weighted aggregation:

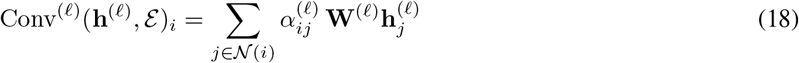

However, this approach can slow down the training process, as the number of nodes involved in the backpropagation scales roughly as *O*(*S*^*L*^) where *S* is the batch size and *L* the number of graph convolution layers. We discuss some cheaper options in Suppl.A.

### 4.6 Interaction scores

One of the main advantages in having an explicit generative model is the possibility of estimating both cross-feature and cell-feature interaction scores. We can define them as:

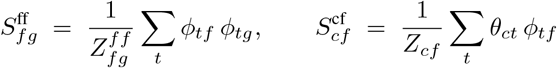

where *θ*_*ct*_ is the proportion of topic *t* in cell *c, φ*_*tf*_ is the loading of feature *f* on topic *t* and 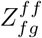 and *Z*_*cf*_ are normalization constants.

### 4.7 Data

Table 1 provides a summary of the data used for the results in Sec.2. The download sources are reported in Sec.5; details on preprocessing are in Suppl.D.

**Table 1:** Summary of the data used. The annotated column refers to whether the annotation came from the download source 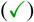, an external source 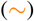 or we performed the annotation via automatic tools 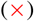. The VisiumHD dataset is made multimodal by considering nucleus RNA and cytoplasmic RNA as two separate modalities only for some models, as discussed in Sec.2.3 and Suppl.D.

| Dataset | Spatial | Multimodal | N. Cells/Spots | Features | Annotated |
| --- | --- | --- | --- | --- | --- |
| TEA-seq (PBMCs) | × | ✓ | 5805 | 2000 RNA + 10000 ATAC + 46 Prot | × |
| ATAC+RNA (Lymphoma) | × | ✓ | 14566 | 2000 RNA + 20000 ATAC | ✓ |
| Spatial Multiome (Mouse Brain) | ✓ | ✓ | 9752 | 25881 RNA + 70470 ATAC | ~ |
| VisiumHD (Colon) P1 | ✓ | ~ | 140066 | 2000 RNA<sub>nucleus</sub> + 2000 RNA<sub>cytoplasm</sub> | ✓ |
| VisiumHD (Colon) P2 | ✓ | ~ | 105622 | 2000 RNA<sub>nucleus</sub> + 2000 RNA<sub>cytoplasm</sub> | ✓ |

## 5 Data availability

The B-Lymphoma dataset was downloaded from the Zenodo repository of SHARE-Topic: https://zenodo.org/records/10418760. The TEA-seq dataset is accessible through GEO, dataset GSE158013, sample GSM5123951, link: https://www.ncbi.nlm.nih.gov/geo/query/acc.cgi?acc=GSE158013. The B-Lymphoma dataset can be downloaded from the 10X Genomics website, at: https://www.10xgenomics.com/datasets/fresh-frozen-lymph-node-with-b-cell-lymphoma-14-k-sorted-nuclei-1-standard-2-0-0.

The spatial RNA+ATAC dataset was downloaded from the Zenodo repository of SpatialGlue: https://zenodo.org/records/10362607. The annotation was downloaded from the COSMOS repository: https://zenodo.org/records/13932144.

The VisiumHD samples can be downloaded at https://www.10xgenomics.com/datasets/visium-hd-cytassist-gene-expression-libraries-of-human-crc and https://www.ncbi.nlm.nih.gov/geo/query/acc.cgi?acc=GSE280318. The annotation can be found at https://github.com/10XGenomics/HumanColonCancer_VisiumHD/tree/main/MetaData.

## 6 Reproducibility

The repository for the package is available at https://github.com/fcaretti/TopOmics. The code to reproduce the results is in the paper folder of the same repository. All the tests were conducted on a single workstation with 112 Intel Xeon Gold 6238R CPU @ 2.20GHz and a NVIDIA A100-PCIE-40GB GPU.

## 7 Acknowledgements

F.C. thanks Magnus Rattray, Syed Murtuza Baker, Sokratia Georgaka, and Esra Busra Isik (University of Manchester) for the many discussions and feedback, and Luca Lasagna (Università di Torino) for the feedback on Sec.2.2. F.C. and G.S. acknowledge the support of the Italian Association for Cancer Research (AIRC) under grant IG 27631. G.S. acknowledges the co-funding from Next Generation EU, in the context of the National Recovery and Resilience Plan, Investment PE1 - Project FAIR “Future Artificial Intelligence Research”. This resource was co-financed by the Next Generation EU [DM 1555 del 11.10.22].

## 8 Authors contribution

F.C., N.E.K. and G.S. conceptualized the work. F.C. implemented the methods, perform the comparisons, produced the figures. F.C. and G.S. wrote the manuscript. All authors revised and approved the manuscript.

## A Other options in the TopOmics library

### Regularization

We optionally include two regularization terms, both set to 0 by default:

- Entropy regularization: *ℒ*_entropy_= *λ*_*H*_ *H*(***θ***) encourages diverse topic usage within cells.
- Topic variance regularization: *ℒ*_var_ = *λ*_*V*_ ∑_*k*_ Var_*n*_(*θ*_*n,k*_) encourages different cells to use different topics, preventing topic collapse.

### Batch effect correction

We provide a simple method for removal of batch effects. Popular amortized inference methods, such as SCVI[6], remove the dependency of the latent representation on given covariates by conditioning the decoder on the set of covariates; in practice, this is done by feeding the decoder with a tensor of covariates stacked on top of the latent representations. Although this method is extremely flexible, it is impossible to control the correction that is applied, and whether some biological effects are being removed during the integration; on the opposite, we can define an explicit parameterization for the set of covariates that we want to control. Specifically, we implement a procedure analogous to [17] in which the batch effect correction is the product of two independent parameters: a gene-batch correction 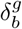 and a topic-level correction 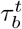.

### Feature Background

To separate a generic expression baseline from topic-specific variation, we follow a practice introduced in [33] and STAMP [17] and add an additive, per-feature background 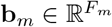 to the topic-feature logits before the softmax. This effect is initialized empirically:

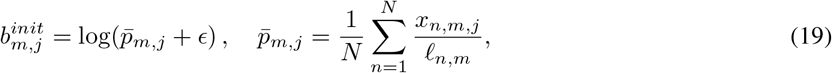

The background is then treated as a latent variable with a Normal prior centered at the initialization value 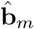,

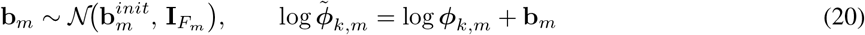

### Multimodal integration

The modality-weighting scheme presented in Sec.4 is far from being the only available one. For example, MultiVI[20] also combines the modality with a MoE strategy, combined with a KL divergence-based loss term that pushes the modalities to generate similar latent representations. The standard Mixture-of-Experts strategy for variational autoencoders, introduced in [30], suffers from lack of expressivity, especially in the cases in which the modalities are significantly unbalanced; however, our training scheme can introduce some problems during generalization, and can therefore be substituted with a simpler version in which the modality weights *ω*_*m*_ are not cell specific but still learnt during training.

### Spatial information aggregation

Although the Graph Attention method explained in Sec.4 is the default TopOmics option, we also provide two popular alternatives already implemented. The first is the swap of the Graph Attention layer in favor of a simpler Graph Convolution one:

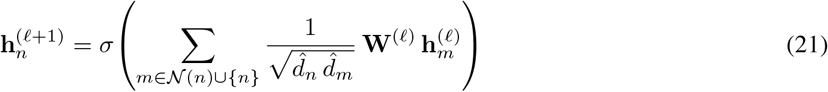

The second option, introduced in [17], is the use of a Simplified Graph Convolution, that is, a simple feature-by-feature average over the features, without trainable parameters.

In practice, this choice leads to pre-computing the input features once at the beginning of training and never updating them, speeding up the training process significantly to the point in which it is practically equivalent to the runtime of variations that do not consider spatial information.

Finally, we also provide as an option a skip connection parameter 0 ≤ *ρ* ≤ 1 that can weight the influence of the neighbors on the latent representation of a node at a given layer:

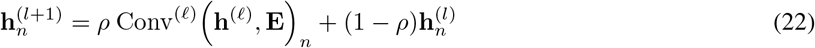

In practice, as the likelihood term of the objective function only considers the single spot reconstruction, this parameter allows the model to only pool the information that is needed to denoise the reconstruction; on the tested datasets, *ρ* ≈ 0.25, leading to worse performances on the tested metrics compared to the settings showed in the main body of the paper.

### Gaussian LDA

The Gaussian variant of the topic model is implemented as a drop-in likelihood within the amortized LDA framework. Each cell *n* and modality *m* is modelled as 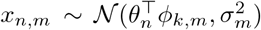, where the topic-feature parameters 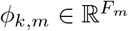 are left unconstrained. Cell-topic proportions *θ*_*n*_ follow the standard Dirichlet distribution, while *ϕ*_*k,m*_ admits either a logistic-normal prior Gaussian modalities skip the library-size log-normalisation applied to count modalities.

Another provided option is the standard Multinomial likelihood, the standard for topic models:

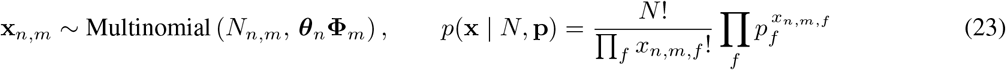

where *N*_*n,m*_ = ∑_*f*_ *x*_*n,m,f*_ is the total count for cell *n* in modality *m*.

### Differences among the prior choices

Fig.7 shows a comparison between the Horseshoe prior with default parameters, and Dirichlet distributions in the case of *α* = 0.05, 1 and 5, where 0.05 is the default in the case of *K* = 20 topics. For lower values of *α*, the prior concentrates the density at the corners of the simplex, generating very sparse topic distributions, while for higher values of *α* the prior cannot generate samples that commit to a single topic. The Horseshoe prior, instead, creates a prior with a “threshold” that, if overcome by the signal, can let the topic proportion grow, hence the higher density stripes, even though the peak of the density is in the middle.

## B 10x Multiome sample of B-Lymphoma lymph node

We show the results of the application of our model to publicly available ATAC-seq sample of lymph node tumor from a patient diagnosed with diffuse small lymphocytic lymphoma. We compare our model to MOFA+[9], MultiVI[20] and SHARE-topic[15]. We evaluate the structure of the latent space using three different metrics: Adjusted Rand Index, Normalized Mutual Information and the accuracy of a kNN classifier trained on 80% of the cells; cell type annotations were provided by the source. Fig.8 shows that the amortized topic model performs comparably to MOFA+ and multiVI while outperforming significantly SHARE-Topic across all metrics. We also notice that the other models are better able to separate T cells from T-cycling cells, while the amortized topic model performs better on the rarer cell types, as observable in the higher balanced accuracy, in which we weight equally all the cell types independently of their relative abundance.

**Figure 7:**
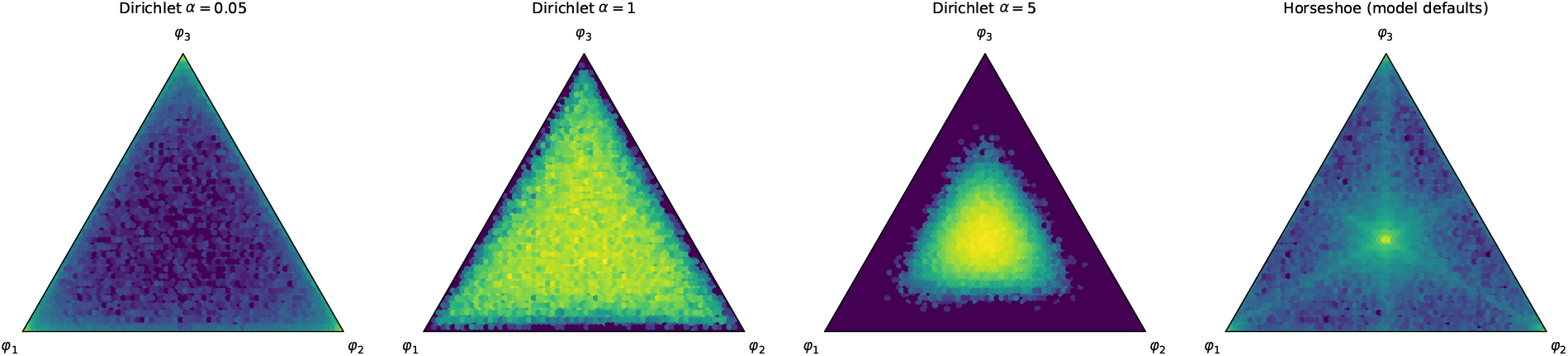
Comparison between the Dirichlet prior distribution with different parameters *α* and the Horseshoe prior, in the case of three features. Color is given by empirical log-density estimated by drawing 50 000 samples

Fig.8f shows the spatial distribution of the topics: in this dataset, almost all the information about the cell type is carried by a single topic, except for the T-cells that are the most abundant type and require the combination of topics 0, 3 and 9 to be characterized.

**Figure 8:**
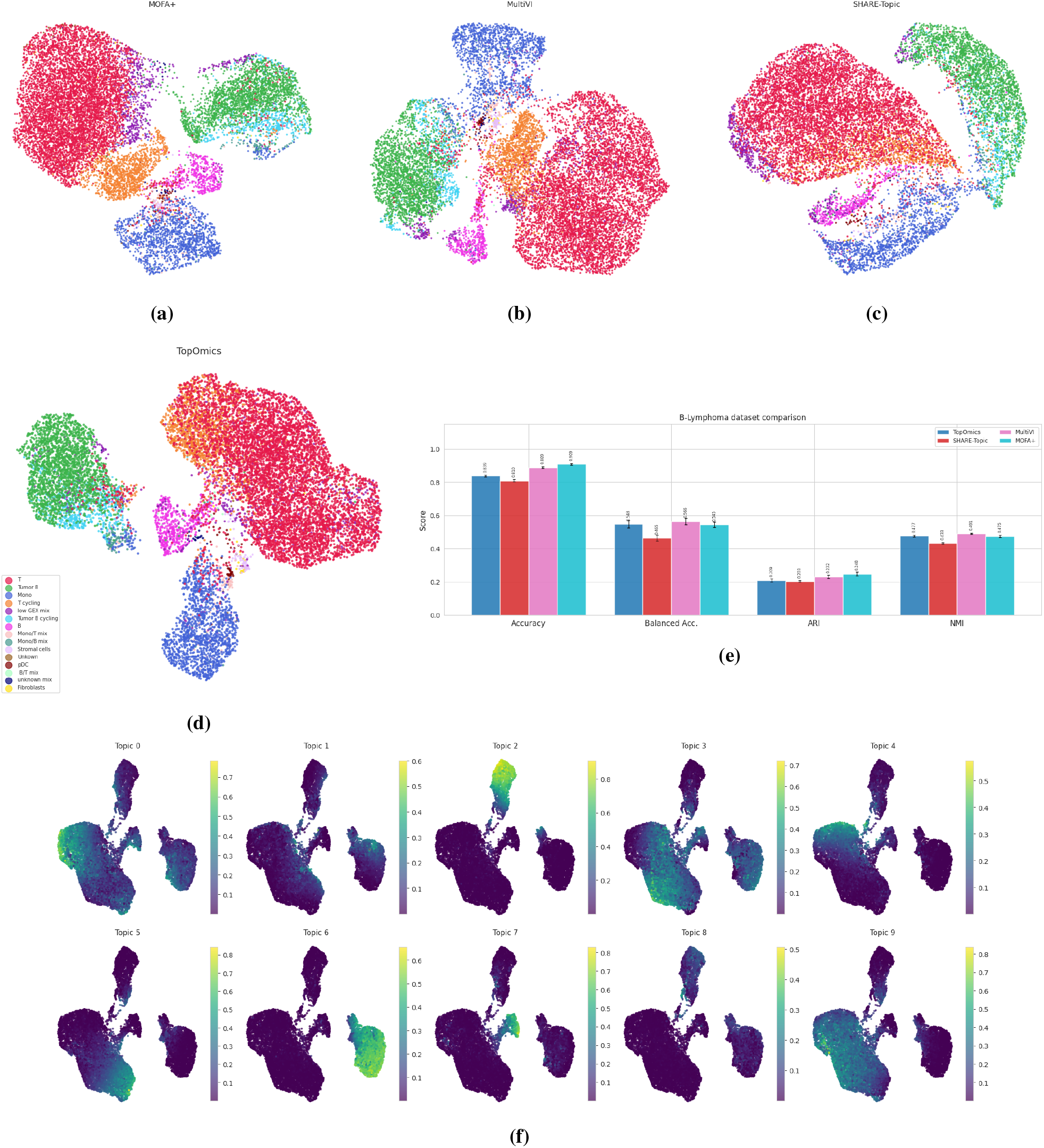
8a8b8c8d: UMAPs of TopOmics latent representation colored by cell type and comparisons with the other tested models. 8e: Accuracy, Accuracy normalized by cell type abundance, ARI (Adjusted Rand Index), and NMI (Normalized Mutual Information) for the 4 tested models. 8f: spatial distribution for every topic

## C Additional plots for the main datasets

Fig.9 shows the distribution of topics using both the available priors on both VisiumHD samples.

**Figure 9:**
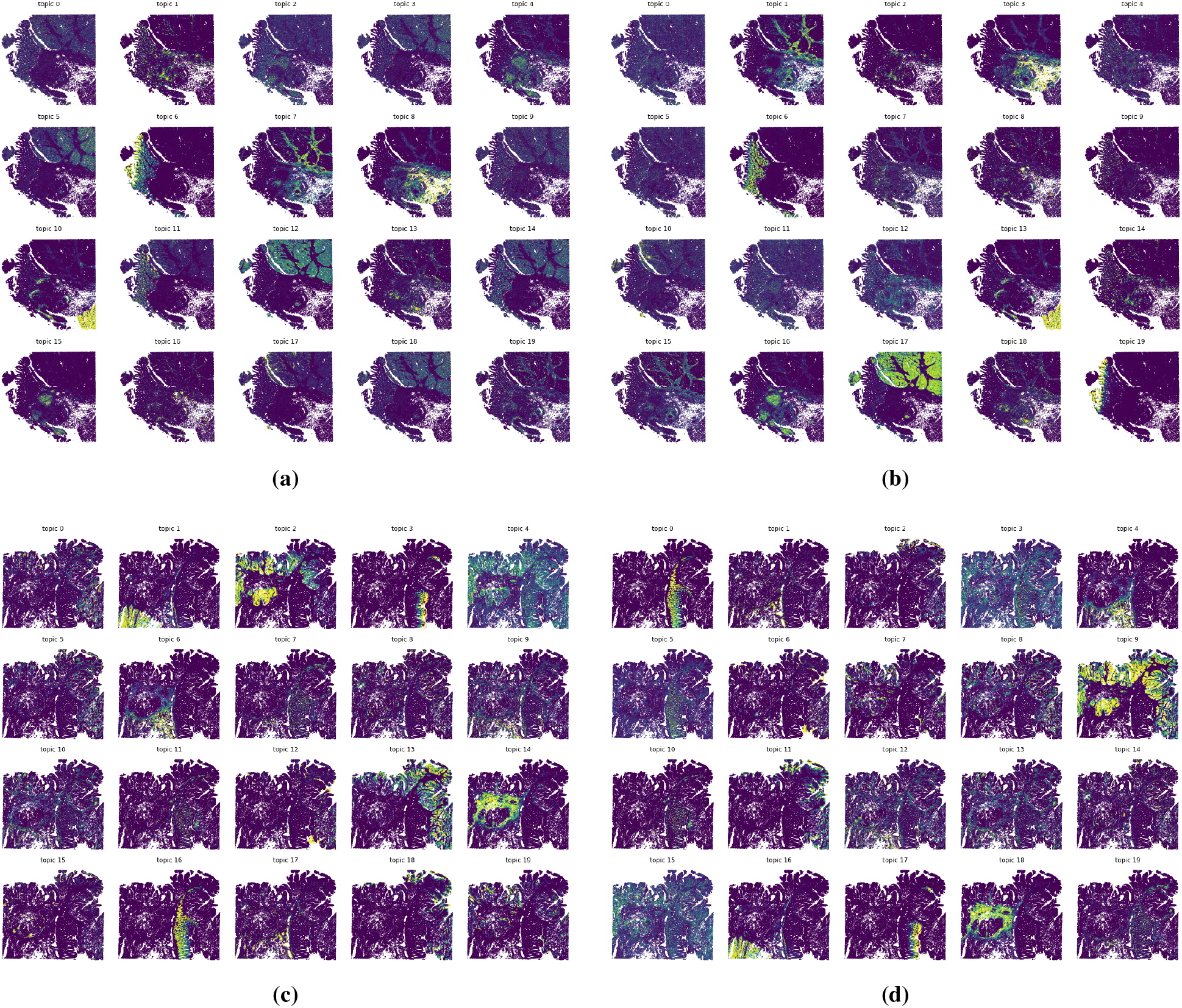
Topic distributions for sample P1 and Dirichlet prior (9a), P1 and Horseshoe prior (9b), P2 and Dirichlet prior (9c) and P2 and Horseshoe prior (9d).

Fig.10 shows examples of the segmentation of cells and nuclei in the VisiumHD P2 sample.

**Figure 10:**
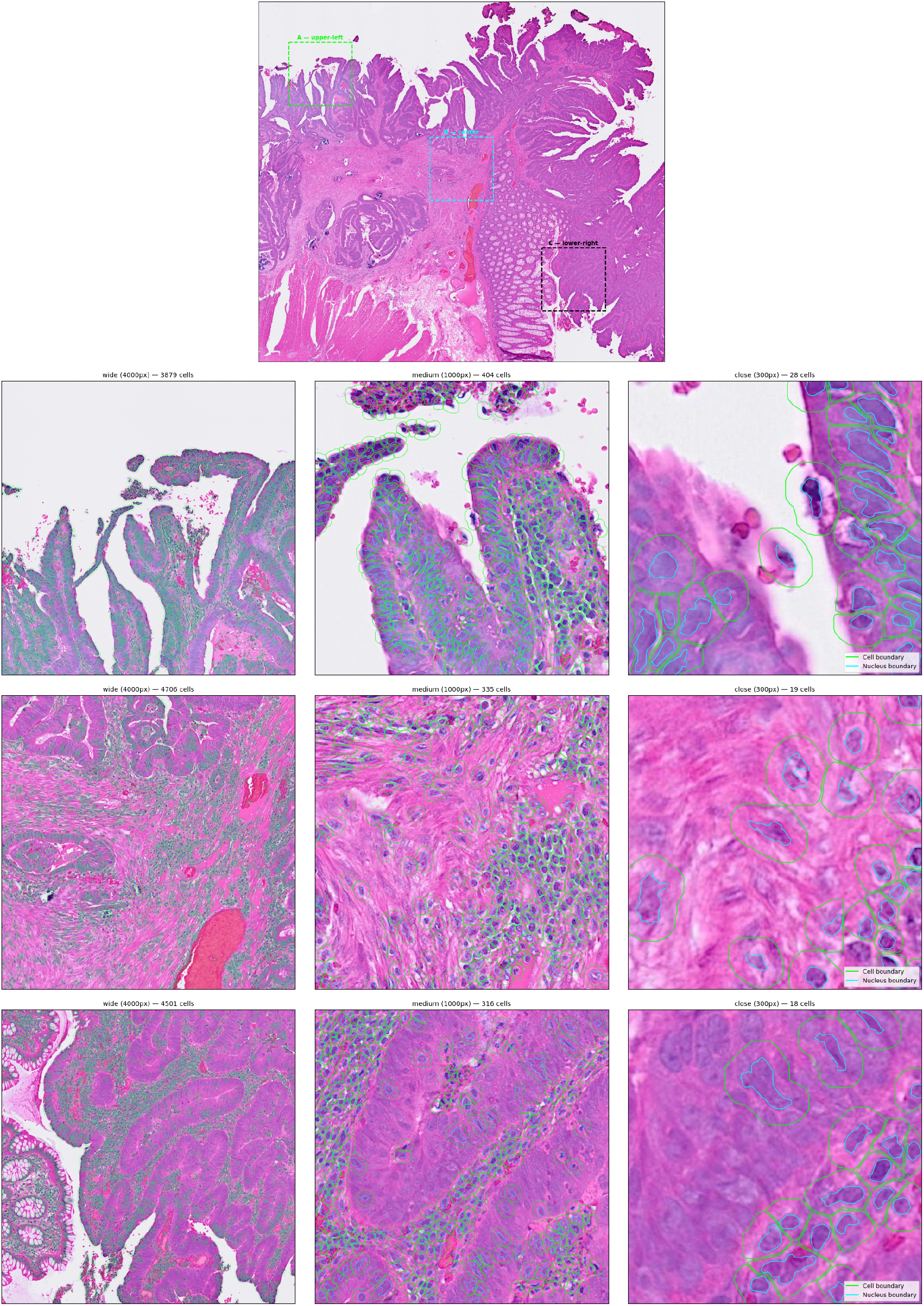
Example visualization of cells in three different areas of the VisiumHD P2 sample

Fig.11 demonstrates that in the TEA-seq analysis, higher cross-interaction scores between transcriptome and open chromatin regions inversely correlate with the distance between the region and the Transcription Starting Site (TSS).

**Figure 11:**
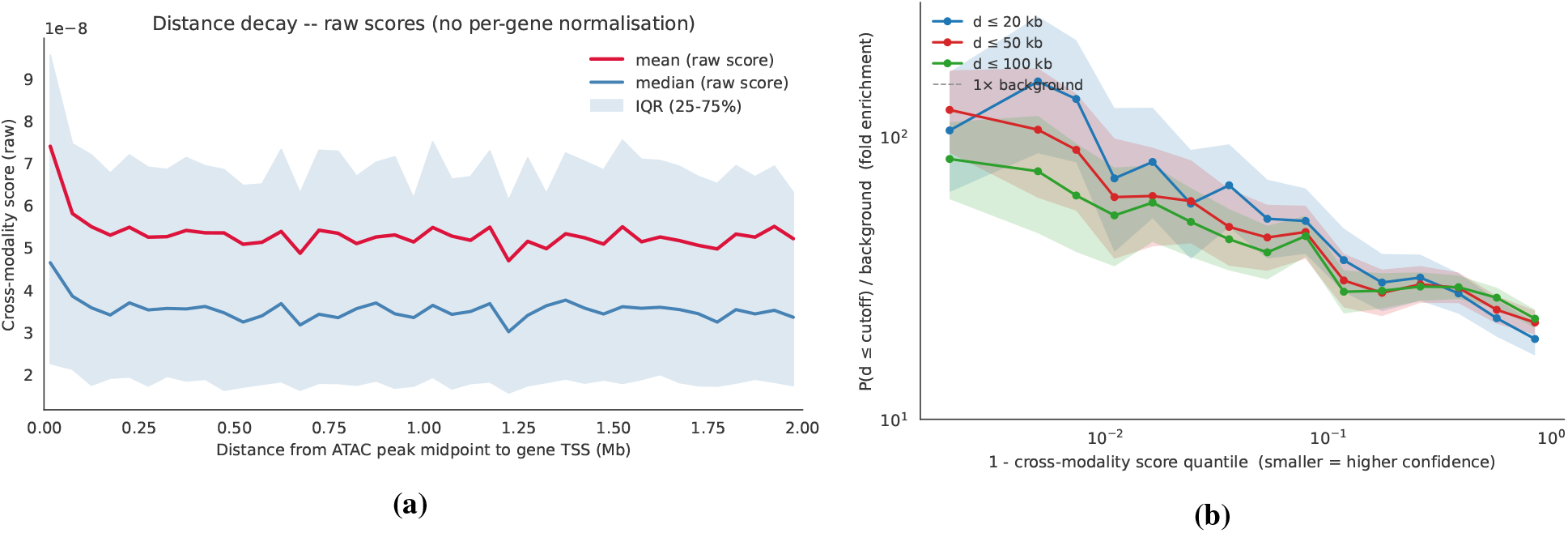
11a: Decay of the average interaction score in the TEA-seq dataset as the distance of the chromatin region and the Transcription Starting Site (TSS) increases. 11a: ratio of distances in a given window per percentile of the cross interaction score

Cross-modality interaction scores were calculated as in Fig.4. In Fig.11a, we plot the interaction score against the distance: it is clearly observable how the main correlation happens between 0 and 25kb of distance. In Fig.11b we first filter the interaction scores to consider only couples closer than 2Mbs, and then divide the interaction scores into quantiles, where the separation is chosen in log-scale to create equal spacing when plotting. Then, we calculate how many gene-region couples in the quantile are closer than 20, 50 or 100 kbs. To allow for a comparison between the different distances, we plot the ratio of this number and the number of couples in the quantile that have a distance higher than 2Mbs, that should not present any meaningful correlation.

## D Figure details

The code to reproduce the results is available at the TopOmics repository under the paperfolder.

Figure 1. We benchmarked the wall-clock training time of four multimodal integration methods (TopOmics, MOFA+, MultiVI, and SHARE-Topic) as a function of dataset size along two axes: number of cells and number of features.

A SHARE-Topic model was first fitted on the full lymphoma RNA+ATAC multiome dataset (comparison of models on the datasets can be found in Suppl.B) using *K*=20 topics, 500 initial burn-in iterations with automatic burn-in detection, and 200 retained posterior samples. Chromatin accessibility counts were binarised before fitting. Synthetic multimodal datasets were then drawn from the fitted posterior, with ATAC counts re-binarised after generation.

We generated synthetic datasets of {1k, 5k, 10k, 50k, 100k, 200k} cells , each retaining the top 2 000 RNA highly variable genes (Seurat v3) and top 20 000 ATAC peaks (selected by variance). Each method was trained 5 times per cell count with *K*=20 latent dimensions/topics, batch size 256, and a maximum of 500 epochs. MultiVI and TopOmics used early stopping (patience 50 for TopOmics; MultiVI defaults). MOFA+ was run in `convergence_mode=“fast”`; SHARE-Topic used 500 burn-in iterations, using an automatic burn-in stopping in the case in which the log-likelihood does not decrease for 50 iterations, and 200 posterior sampling iterations. MOFA+ and SHARE-Topic were trained up until it was allowed by our resources, reported in Sec.6.

Figure 2. We evaluated TopOmics against four multimodal integration baselines(MOFA+, MultiVI, SpatialGlue, and COSMOS) on the Mouse Brain spatial multi-omics dataset from [8], which pairs spatial RNA-seq with spatial ATAC-seq on the same tissue section.

We downloaded RNA and ATAC count matrices from the SpatialGlue data release. ATAC counts were binarised. A spatial kNN graph (*k*=4, Euclidean distance on spatial coordinates) was constructed and shared across modalities. Ground-truth layer annotations (8 brain areas, excluding “others”) were obtained from the COSMOS data release.

All methods used a latent dimensionality of *K*=20 for the clustering panels and *K*=10 for the latent factors spatial distribution images for easier visualization. MOFA+ was trained on log-normalised, scaled RNA and ATAC (`convergence_mode=“medium”`). MultiVI was trained on raw counts with 80/20 train/validation split and early stopping (max 300 epochs, *n*_hidden_=128). SpatialGlue used PCA (50 components) on log-normalised RNA and LSI (50 components) on binarised ATAC as input features, and was trained for 500 epochs (learning rate 10^*−*4^ , weight factors [1, 5, 1, 1]). COSMOS used 3 000 HVGs (Seurat v3, normalised, log-transformed, scaled) for RNA and 50 LSI components for ATAC. It was trained with spatial regularisation strength 0.05 (default) and 0.1, *z*_dim_=*K*, learning rate 10^*−*3^, WNN pre-training for 500 epochs, total training up to 1 000 epochs with early stopping (patience 10 before, 30 after regularisation onset, minimum 200 epochs). We also tested other values of the regularization strength (0.0 and up to 0.2) and reported results only for the default value and for the value with the best performance metrics.

TopOmics (`MultimodalAmortizedLDA`) was trained on raw RNA counts (Gamma-Poisson likelihood) and binarised ATAC peaks (Bernoulli likelihood), with per-cell modality weights (`weight_mode=“cell”`), mixture-of-experts aggregation, a 2-layer GATv2 spatial encoder (initial skip-connection *α*=0.2, learned during training, *k*=5 spatial neighbors), and 1*/K* symmetric Dirichlet prior on cell-topic proportions. Two feature-prior variants were compared: Dirichlet (logistic-normal) and Horseshoe. Models were trained with learning rate 10^*−*2^, batch size 256, and 80/20 train/validation split.

For the spatial clustering panels, *k*-means with *k*=15 clusters was applied to each method’s 20D latent representation, based on the Euclidean distance for all models except TopOmics (Hellinger distance). Quantitative evaluation used bootstrapped (*n*=50 resamples) Accuracy and NMI against the 8 ground-truth brain areas, computed at both *k*=8 (matching the number of annotated classes) and *k*=15. Spatial coherence was assessed via mean Moran’s *I* across latent dimensions, computed on the spatial neighbor graph.

Figure 3. We applied TopOmics to a Visium HD Human Colon Cancer FFPE dataset (10x Genomics), exploiting the sub-cellular resolution to treat nucleus and cytoplasm RNA as two separate modalities.

Cell and nucleus boundaries were segmented from the H&E image using QuPath, producing a GeoJSON with paired cell and nucleus polygons. Cytoplasm polygons were obtained by subtracting the nucleus from the cell polygon (pairs where the residual area was *<* 5 px^2^ were discarded). Each Visium HD 2 *µ*m bin was assigned to a nucleus or cytoplasm compartment via spatial join, resulting in two AnnData matrices (cells × genes): one for nucleus RNA and one for cytoplasm RNA. Cells present in both compartments were retained.

The top 2000 highly variable genes were selected once on the *combined* (nucleus + cytoplasm) count matrix using Scanpy (Seurat v3 flavour), and all models were trained on the same gene sets.

TopOmics (`MultimodalAmortizedLDA`) was set up as a bimodal model with nucleus and cytoplasm as separate modalities, both using a negative binomial likelihood. We tested both the possible topic-feature prior choices. All variants used *K*=20 topics, *n*_hidden_=128, mixture-of-experts aggregation, batch size 128, 90/10 train/validation split, and up to 200 epochs. A symmetric kNN graph (*k*=10 built on the Euclidean distance of cell centroids was used to inform the graph convolution.

scVI was trained on either nucleus or cytoplasm only counts or the summed nucleus+cytoplasm counts (*n*_latent_=20, *n*_hidden_=128, negative-binomial likelihood, max 200 epochs, batch size 128, early stopping with patience 15, 90/10 split). COSMOS was trained on the (nucleus, cytoplasm) pair after independent per-modality preprocessing (normalizing the total to 10^4^, log 1*p*, and features rescaling). The model was trained with *z*_dim_=20, learning rate 10^*−*3^, with a spatial-feature graph of *k*=5 neighbors, 100 total epochs (with the WNN-fusion phase activated at epoch 50), spatial regularization strength 0.1.

Bootstrapped (*n*=10) MiniBatch *k*-means clustering (with *k* set to the number of ground-truth classes) was evaluated against each annotation via ARI, NMI, and majority-vote cluster accuracy. Spatial coherence was measured as the mean Moran’s *I* across latent dimensions, computed on the spatial *k*NN graph.

Figure 4. We evaluated TopOmics on the TEA-seq dataset, which simultaneously profiles RNA, ATAC, and surface protein in human peripheral blood mononuclear cells (PBMCs).

The MuData object (5 805 cells) contains three modalities: RNA, ATAC, and protein (46 ADTs). ATAC counts were binarised. RNA was filtered to the top 2 000 highly variable genes (Seurat v3) and ATAC to the top 10 000 highly variable peaks; all 46 protein features were retained. Cell type annotations were obtained by running CellTypist[26] with the `Immune_All_Low` reference model and majority voting, resulting in 10 cell types.

MultiVI was trained with Negative Binomial likelihood function for RNA and protein counts and Bernoulli for the binarized ATAC counts (*n*_latent_=10, max 300 epochs, early stopping). MOFA+ was trained on log-normalised, scaled RNA and ATAC. Single-modality AmortizedLDA baselines (one per modality) were also trained with *K*=10 topics.

TopOmics was trained with *K*=10 topics, logistic-normal feature prior, per-cell modality weights (`weight_mode=“cell”`) with mixture-of-experts aggregation, per-gene learnable dispersion, Gamma-Poisson likelihood for RNA and protein, Bernoulli likelihood for ATAC, 1*/K* symmetric Dirichlet cell-topic prior, batch size 128, 80/20 train/validation split, and up to 500 training epochs.

All methods were evaluated with bootstrapped (*n*=50 resamples) *k*-nearest-neighbour classification (*k*=5, 80/20 stratified split, accuracy and balanced accuracy), as well as *k*-means clustering (ARI, NMI) at *k* equal to the number of annotated cell types. For TopOmics and single-modality AmortizedLDA, Hellinger distances were considered instead of Euclidean distances.

In Fig. 4i, the top 8 marker features per cell type were selected for each modality using a topic-weighted specificity score: the mean topic proportions per cell type were used to weight the feature-topic distribution *ϕ*, and features with the highest cluster-specific score (normalised by total score across clusters) were retained. Expression values were *z*-scored and clipped to [−2, 2]. Cells were ordered by cell type and, within each type, by dominant topic assignment.

Figs.4j4k4l show feature-feature interactions for each combination of two modalities. We started by clustering the cells according to their proteins profile, in a number of clusters equal to the number of cell types. Then, for both RNA and chromatin we selected the top 5 features by average correlation with each of the clusters. Finally, in each plot, we filter the features (mostly the proteins) by keeping only those whose z-score peak across either row or column exceeds 1.5 at least once, and then order the features on both row and column, by applying Ward hierarchical clustering first and then ordering row and column in descending order. Each feature was colored by the color corresponding to the cell type they are the most characteristic of, calculated through the cell-feature score using only the cells belonging to the cell type.

## E Tables of quantitative results

**Table 2:**
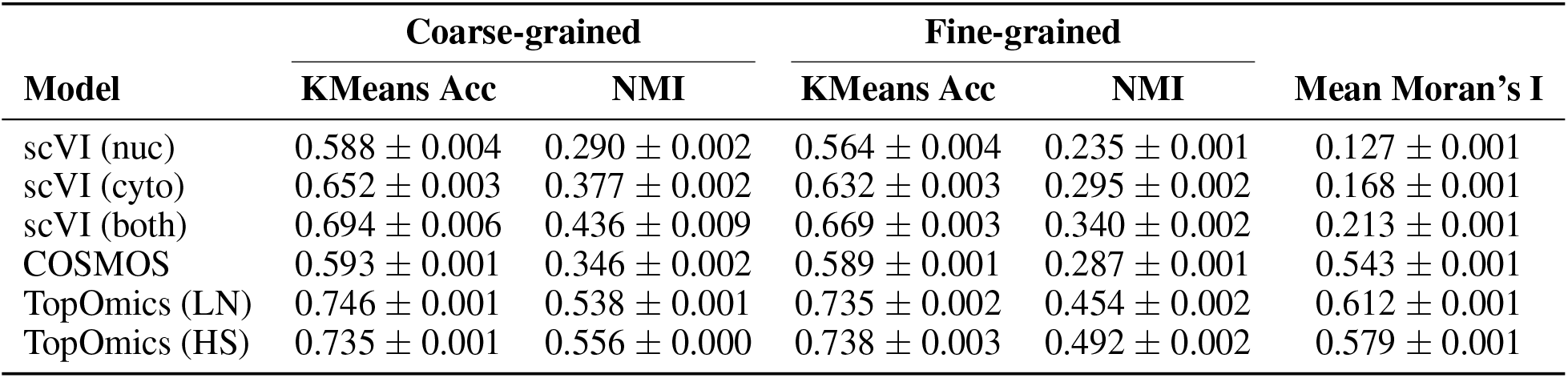
Table referring to metrics for the VisiumHD colorectal cancer dataset shown in Fig.3g.

| Model | Coarse-grained |  | Fine-grained |  | Mean Moran's I |
| --- | --- | --- | --- | --- | --- |
|  | KMeans Acc | NMI | KMeans Acc | NMI |  |
| scVI (nuc) | 0.588 ± 0.004 | 0.290 ± 0.002 | 0.564 ± 0.004 | 0.235 ± 0.001 | 0.127 ± 0.001 |
| scVI (cyto) | 0.652 ± 0.003 | 0.377 ± 0.002 | 0.632 ± 0.003 | 0.295 ± 0.002 | 0.168 ± 0.001 |
| scVI (both) | 0.694 ± 0.006 | 0.436 ± 0.009 | 0.669 ± 0.003 | 0.340 ± 0.002 | 0.213 ± 0.001 |
| COSMOS | 0.593 ± 0.001 | 0.346 ± 0.002 | 0.589 ± 0.001 | 0.287 ± 0.001 | 0.543 ± 0.001 |
| TopOmics (LN) | 0.746 ± 0.001 | 0.538 ± 0.001 | 0.735 ± 0.002 | 0.454 ± 0.002 | 0.612 ± 0.001 |
| TopOmics (HS) | 0.735 ± 0.001 | 0.556 ± 0.000 | 0.738 ± 0.003 | 0.492 ± 0.002 | 0.579 ± 0.001 |

**Table 3:**
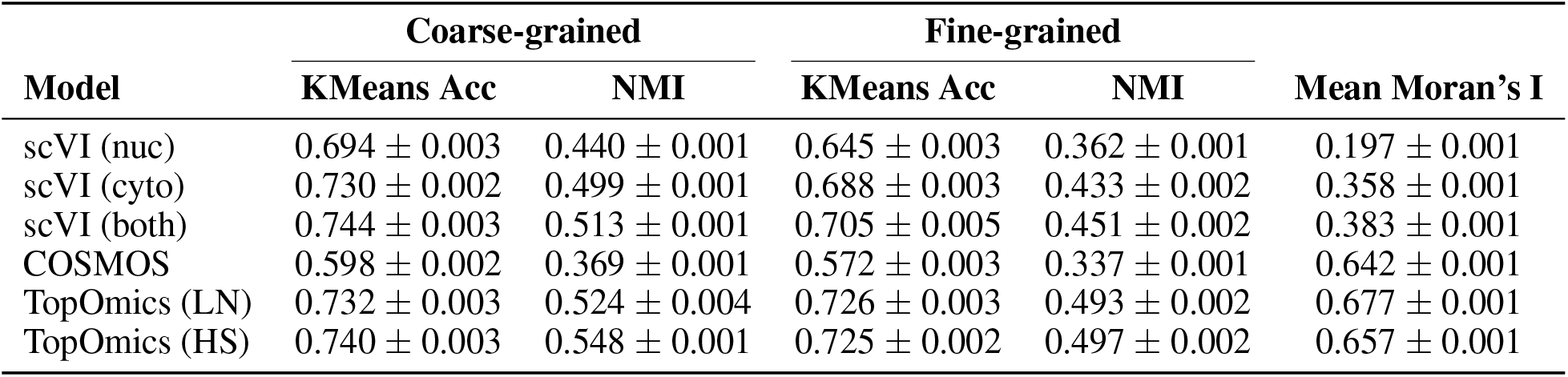
Table referring to metrics for the VisiumHD colorectal cancer dataset shown in Fig.3h.

| Model | Coarse-grained |  | Fine-grained |  | Mean Moran's I |
| --- | --- | --- | --- | --- | --- |
|  | KMeans Acc | NMI | KMeans Acc | NMI |  |
| scVI (nuc) | 0.694 ± 0.003 | 0.440 ± 0.001 | 0.645 ± 0.003 | 0.362 ± 0.001 | 0.197 ± 0.001 |
| scVI (cyto) | 0.730 ± 0.002 | 0.499 ± 0.001 | 0.688 ± 0.003 | 0.433 ± 0.002 | 0.358 ± 0.001 |
| scVI (both) | 0.744 ± 0.003 | 0.513 ± 0.001 | 0.705 ± 0.005 | 0.451 ± 0.002 | 0.383 ± 0.001 |
| COSMOS | 0.598 ± 0.002 | 0.369 ± 0.001 | 0.572 ± 0.003 | 0.337 ± 0.001 | 0.642 ± 0.001 |
| TopOmics (LN) | 0.732 ± 0.003 | 0.524 ± 0.004 | 0.726 ± 0.003 | 0.493 ± 0.002 | 0.677 ± 0.001 |
| TopOmics (HS) | 0.740 ± 0.003 | 0.548 ± 0.001 | 0.725 ± 0.002 | 0.497 ± 0.002 | 0.657 ± 0.001 |

**Table 4:**
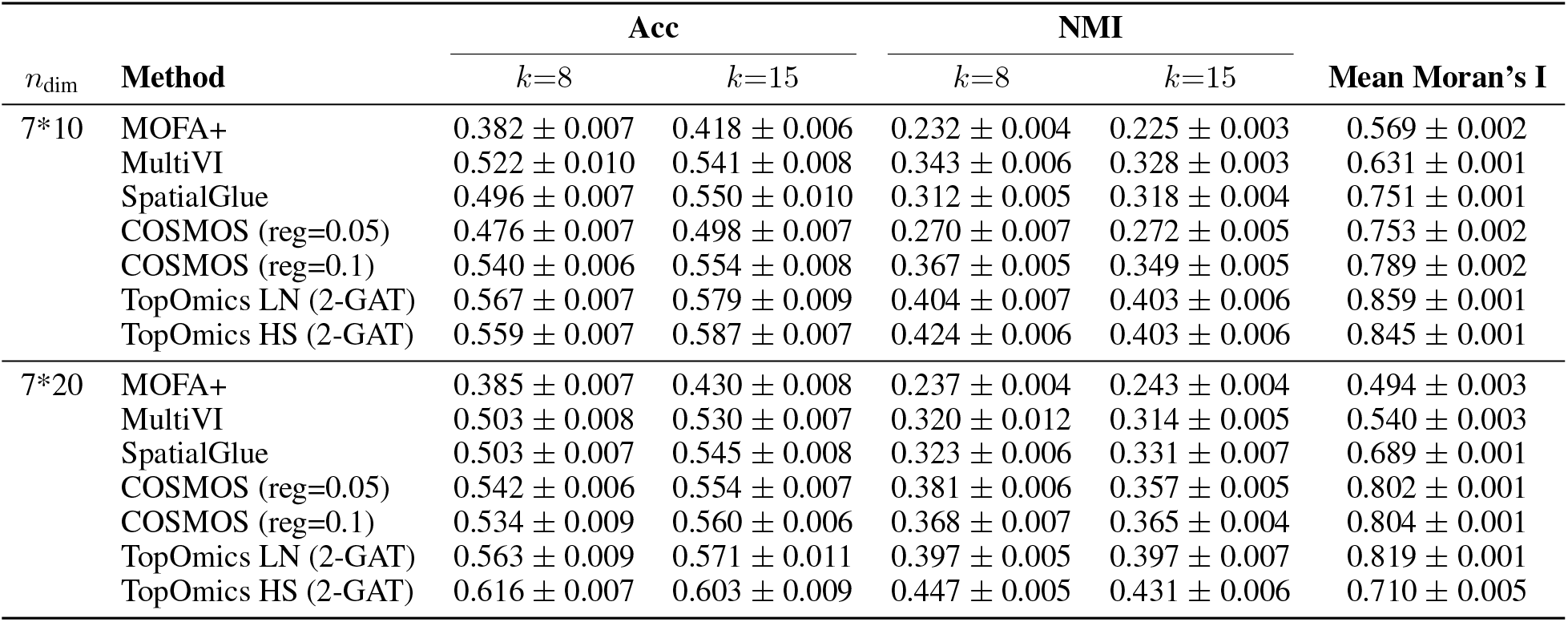
Table referring to metrics for the spatial RNA+ATAC mouse brain dataset shown in Fig.2h.

| $n_{\text{dim}}$ | Method | Acc | | NMI | | Mean Moran's I |
| --- | --- | --- | --- | --- | --- | --- |
| | | $k=8$ | $k=15$ | $k=8$ | $k=15$ | |
| 7*10 | MOFA+ | 0.382 ± 0.007 | 0.418 ± 0.006 | 0.232 ± 0.004 | 0.225 ± 0.003 | 0.569 ± 0.002 |
|  | MultiVI | 0.522 ± 0.010 | 0.541 ± 0.008 | 0.343 ± 0.006 | 0.328 ± 0.003 | 0.631 ± 0.001 |
|  | SpatialGlue | 0.496 ± 0.007 | 0.550 ± 0.010 | 0.312 ± 0.005 | 0.318 ± 0.004 | 0.751 ± 0.001 |
|  | COSMOS (reg=0.05) | 0.476 ± 0.007 | 0.498 ± 0.007 | 0.270 ± 0.007 | 0.272 ± 0.005 | 0.753 ± 0.002 |
|  | COSMOS (reg=0.1) | 0.540 ± 0.006 | 0.554 ± 0.008 | 0.367 ± 0.005 | 0.349 ± 0.005 | 0.789 ± 0.002 |
|  | TopOmics LN (2-GAT) | 0.567 ± 0.007 | 0.579 ± 0.009 | 0.404 ± 0.007 | 0.403 ± 0.006 | 0.859 ± 0.001 |
|  | TopOmics HS (2-GAT) | 0.559 ± 0.007 | 0.587 ± 0.007 | 0.424 ± 0.006 | 0.403 ± 0.006 | 0.845 ± 0.001 |
| 7*20 | MOFA+ | 0.385 ± 0.007 | 0.430 ± 0.008 | 0.237 ± 0.004 | 0.243 ± 0.004 | 0.494 ± 0.003 |
|  | MultiVI | 0.503 ± 0.008 | 0.530 ± 0.007 | 0.320 ± 0.012 | 0.314 ± 0.005 | 0.540 ± 0.003 |
|  | SpatialGlue | 0.503 ± 0.007 | 0.545 ± 0.008 | 0.323 ± 0.006 | 0.331 ± 0.007 | 0.689 ± 0.001 |
|  | COSMOS (reg=0.05) | 0.542 ± 0.006 | 0.554 ± 0.007 | 0.381 ± 0.006 | 0.357 ± 0.005 | 0.802 ± 0.001 |
|  | COSMOS (reg=0.1) | 0.534 ± 0.009 | 0.560 ± 0.006 | 0.368 ± 0.007 | 0.365 ± 0.004 | 0.804 ± 0.001 |
|  | TopOmics LN (2-GAT) | 0.563 ± 0.009 | 0.571 ± 0.011 | 0.397 ± 0.005 | 0.397 ± 0.007 | 0.819 ± 0.001 |
|  | TopOmics HS (2-GAT) | 0.616 ± 0.007 | 0.603 ± 0.009 | 0.447 ± 0.005 | 0.431 ± 0.006 | 0.710 ± 0.005 |

**Table 5:** Table referring to metrics for the TEA-seq dataset shown in Fig.4g.

| Method | Acc | bAcc | ARI | NMI |
| --- | --- | --- | --- | --- |
| TopOmics | 0.913 ± 0.005 | 0.775 ± 0.031 | 0.395 ± 0.016 | 0.647 ± 0.006 |
| MultiVI | 0.911 ± 0.007 | 0.791 ± 0.033 | 0.433 ± 0.015 | 0.668 ± 0.007 |
| MOFA+ | 0.918 ± 0.006 | 0.740 ± 0.038 | 0.329 ± 0.022 | 0.562 ± 0.010 |
| AmortizedLDA (RNA) | 0.846 ± 0.007 | 0.616 ± 0.029 | 0.311 ± 0.007 | 0.547 ± 0.005 |
| AmortizedLDA (ATAC) | 0.793 ± 0.009 | 0.403 ± 0.016 | 0.281 ± 0.011 | 0.446 ± 0.007 |
| AmortizedLDA (Protein) | 0.903 ± 0.007 | 0.727 ± 0.026 | 0.367 ± 0.020 | 0.616 ± 0.008 |

## F KL calculation

Here we derive explicitly the KL term for the cell-topic distribution. We recall that

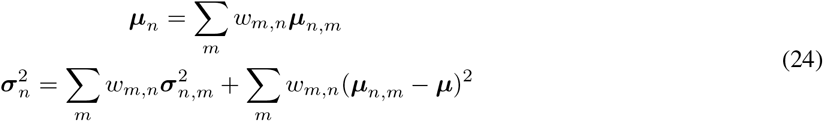

Without loss of generality, we restrict our analysis to the 2-modalities case. We take the default option for the modality weights:

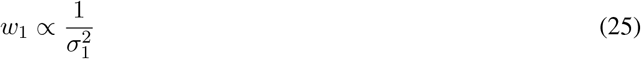

and in particular, for normalization purpose:

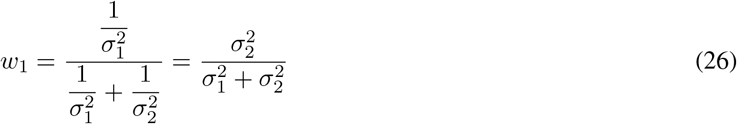

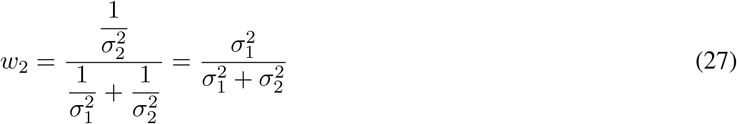

Then, the aggregation of both modalities leads to a Gaussian distribution (before the SoftMax function) with parameters:

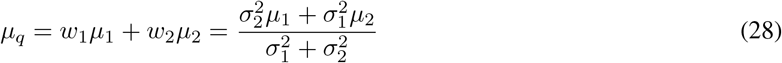

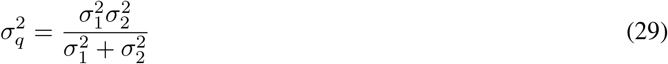

We notice that by choosing *α*_*k*_ = *α* and substituting into Eqs. **??** and **??** we get:

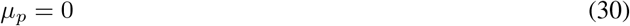

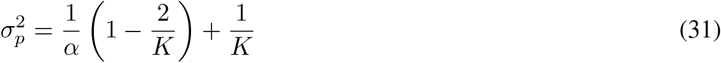

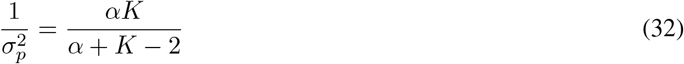

Substituting every term in the KL divergence yields the following:

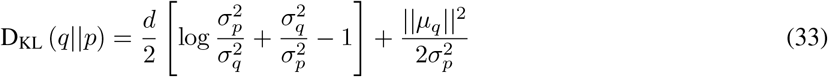

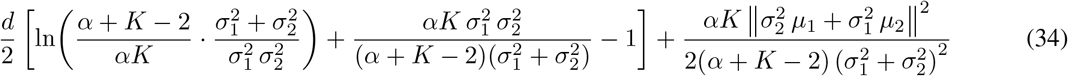

Focusing on the last term, we notice that the KL divergence is minimized if *µ*_1_ = *µ*_2_ = 0 and if *µ*_1_ and *µ*_2_ point to opposite directions (this is true in general, independently of the specific choice of weights).

The choice of the Mixture-of-Experts therefore slightly pushes the modalities to encode similar representations.

